# Imputation-free reconstructions of three-dimensional chromosome architectures in human diploid single-cells using allele-specified contacts

**DOI:** 10.1101/2021.10.04.462972

**Authors:** Yoshito Hirata, Arisa H. Oda, Chie Motono, Masanori Shiro, Kunihiro Ohta

## Abstract

The sparseness of chromosomal contact information and the presence of homologous chromosomes with very similar nucleotide sequences make Hi-C analysis difficult. We propose a new algorithm using allele-specific single-nucleotide variations (SNVs) to reconstruct the three-dimensional (3D) chromosomal architectures from the Hi-C dataset of single diploid cells. Our algorithm has a function to discriminate SNVs specifically found between homologous chromosomes to our “recurrence plot”-based algorithm to estimate the 3D chromosome structure, which does not require imputation for ambiguous segment information. The new algorithm can efficiently reconstruct 3D chromosomal structures in single human diploid cells by employing only Hi-C segment pairs containing allele-specific SNVs. The datasets of the remaining pairs of segments without allele-specific SNVs are used to validate the estimated chromosome structure. This approach was used to reconstruct the 3D structures of human chromosomes in single diploid cells at a 1-Mb resolution. Introducing a subsequent mathematical measure further improved the resolution to 40-kb or 100-kb. The reconstruction data reveals that human chromosomes form chromosomal territories and take fractal structures where the mean dimension is a non-integer value. We also validate our approach by estimating 3D protein/polymer structures.

## 1. Introduction

The three-dimensional (3D) chromosomal structure plays important roles in various biological processes such as DNA replication and gene regulation. There are two major methods to investigate the 3D chromosomal structures: 1) microscopy-based fluorescent in situ hybridization and 2) chromosome conformation capture techniques combined with deep sequencing and a computational reconstruction (Hi-C) (Le Dily *et al.*, 2017). Recently, the Hi-C method has been applied to individual human cells (Nagano *et al.*, 2013; Stevens *et al.*, 2017; Flyamer *et al.*, 2017). Although several methods have been developed to reconstruct the 3D chromosomal structure in human haploid cells (Paulsen *et al.*, 2015; Carstens *et al.*, 2016; Hirata *et al.*, 2016), the available methods for diploid cells are mostly for ensemble Hi-C data (Cauer *et al.*, 2019).

Among them, Carstens *et al.* (2016) has been used for a single diploid cell. They combined ambiguous distance constraints with the inverse sixth powers of distances to realize the “OR” operation, or the circumstance where chromosome segments of a paternal or maternal allele satisfy some distance constraints. They claim that bias can be avoided in assigning alleles for each contact. However, their results are validated only from the viewpoint of consistency with preexisting results.

A previous report demonstrated an experimental method and its accompanying computational method called imputation, which were proposed to overcome the sparseness of the Hi-C dataset of single diploid cells (Tan *et al.*, 2018). They distinguished two alleles on each homolog by differently labeled single nucleotide variations (SNVs). Then they imputed unlabeled alleles using the information of neighbors by assuming that different alleles typically contact different chromosomal segments. Specifically, they made the following assumptions: (i) two alleles are not close to each other and (ii) alleles do not have similar shapes (Tan *et al.*, 2018). Lastly, they used the imputed allele labels to reconstruct the 3D chromosomal structure with simulated annealing. Since the frequency of SNVs should be insufficient to mark all the sequence segments read from different alleles, only a few percent of segments contain enough information to identify the derived allele. The rest contain only ambiguous information (Supplementary Table 1). Imputation tries to employ this ambiguous segment information to obtain a high-resolution reconstruction. This Hi-C reconstruction algorithm for a single diploid cell is quite powerful, but such imputations may contain above-mentioned assertions that cannot be verified directly.

Here, we propose an alternative imputation-free computational method to reconstruct the 3D structure from Hi-C data for a single diploid cell. This method is an extension of our previous recurrence plot-based reconstruction method (Hirata *et al.*, 2016). The key feature of our method is that only consecutive chromosome segments are assumed to be neighboring. We use the parts of pairs where both alleles on homologs contain SNVs. The remaining pairs of chromosome segments, which have at least one allele without SNVs, were used for the self-validation of the estimated 3D chromosomal structures. Finally, we discuss the validity for the reconstructed 3D structure by checking the similarity and difference between 3D structures for allele pairs.

## 2. Method

### 2.1 Recurrence plot-based reconstructions

We use the similarity between single-cell Hi-C data and a recurrence plot (Eckmann *et al.*, 1987; Marwan *et al.*, 2007) to reconstruct the 3D structure for corresponding chromosomes (Hirata *et al.*, 2016). We apply the same strategy to reproduce the 3D structure from single-cell Hi-C data. However, the present study has two differences compared with Hirata *et al.* (2016). First, only segment pairs containing SNVs are used to calculate local distances between segment pairs. Second, this study initially obtains a coarse reconstruction using the method of Hirata *et al.* (2016), which is subsequently refined by employing the idea of time series forecasting (Sugihara and May, 1990).

### 2.2 Discrimination of homologous chromosomes using data of paired segments with allele-specific SNVs

This section describes the differentiation of two homologous chromosomes for a single diploid cell in the proposed method.

For an ensemble of diploid cells, a Bayesian technique can be used to differentiate maternal alleles from paternal alleles (Carstens *et al.*, 2016). For the single diploid cell data presented in Tan *et al.* (2018), we experimentally used the SNVs to differentiate one allele from the other as a potential genetic marker from the sequencing information. Thus, we focus on pairs of chromosome segments, which are spatially close enough to be detected by the Hi-C experiment and contain SNVs. Only these pairs of chromosome segments (phased pairs) are used. We apply the above algorithm to reproduce the 3D structure for the chromosomes (Fig. 1).

**Figure 1:**
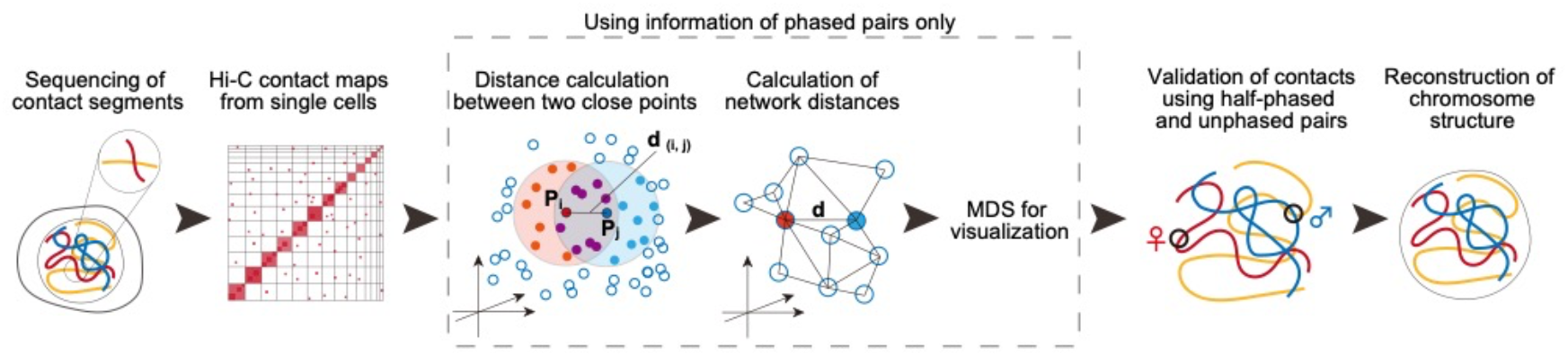
Graphic summary of the reconstruction of the 3D structure for chromosomes given single-cell Hi-C data using the recurrence plot-based method.

Thus, we can use pairs of spatially close chromosome segments where at least one of them may not contain the sequence of SNVs to verify the reconstructed 3D structure (Supplementary Fig. 1). Namely, if two chromosome segments are detected as neighbors, one of the chromosome segments can be identified by its original allele while the origin of the other allele is unknown (half-phased pairs). For half-phased pairs, the distance of the identified segment to the corresponding location of one of two allele segments should be close. If two chromosome segments are detected as neighbors, and neither contains SNVs (unphased pairs), it is impossible to tell which alleles they are from, but the closest distance among four possible pairs between the corresponding allele segments should be close. We will validate this tendency in Section 3.2. Therefore, the novelty of the current work is (i) it applies the method of Hirata *et al.*, (2016) only on phased pairs of single diploid cell Hi-C data, (ii) the reconstruction is refined, and (iii) half-phased and unphased pairs validate the reconstruction.

### 2.3 Summary of the proposed method

The computational procedure of our reconstruction can be summarized as follows. First, we declare that consecutive chromosome segments on the same allele are neighbors (Tanio *et al.*, 2009; Hirata *et al.*, 2016) and neighbors are defined by phased pairs. Second, we construct a network from the Hi-C map (Hirata *et al.*, 2008, 2015, 2016), where each node corresponds to a chromosome segment and each contact corresponds to an edge. Then, we assign a local distance to each edge. The local distance between two chromosome segments can be determined by the ratio of the unshared neighbors against the union of the neighbors at their corresponding rows (Hirata *et al.*, 2008, 2016). Third, we obtain the shortest distances between every pair of nodes (Hirata *et al.*, 2008, 2016). The shortest distances can be regarded as the global distances (Tenenbaum *et al.*, 2000). Fourth, we convert the global distances into point arrangements in 3D space while preserving the distances via multidimensional scaling (Gower, 1966). These point arrangements correspond to our coarse 3D reconstruction at a 1-Mb resolution. Lastly, the four closest neighbors are found for each point at a 40-kb or 100-kb resolution based on the similarity of the connected nodes. Then the weighted averages (Sugihara and May, 1990) of the neighbors’ coarse 3D reconstruction at a 1-Mb resolution are used for a finer reconstruction at a 40-kb or 100-kb resolution. If the last step is removed, then the proposed method coincides with our previous method (Hirata *et al.*, 2016). In addition, if all the local distances are approximated by 1 at the second step, our method agrees with the single-cell Hi-C implementation of Lesne *et al.* (2014). In Section 2.4, we will compare the proposed method with that of Lesne *et al.* (2014). It should be noted that Paulsen *et al.* (2015) defined local distances as a constant for single-cell Hi-C data, but they employed a manifold based learning technique instead of the shortest distance approach used here.

Figure 1 shows a graphic summary of how we reconstructed the 3D structure of the chromosomes from single-cell Hi-C data. In addition, the Supplementary Material contains the mathematical details for the above calculations.

### 2.4 Validation using protein/polymer models

First, we tested the proposed algorithm before the refinement with protein data, which are polymers of amino acids. We used the data of 14 proteins, which are given by the distances between every pair of points. The test protein set was constructed from the CASP13 targets from https://www.predictioncenter.org/download_area/CASP13/targets/, with the official domain definitions (Kryshtafovych *et al.*, 2019; Kinch *et al.*, 2019). We chose all proteins classified as “Free Modeling (FM)” with structures deposited at the Worldwide Protein Data Bank as of March 2020. The length of the resulting proteins ranged between 72–374 amino acids. Supplementary Table 2 lists the proteins used in this study. Two residues are in contact if the distance between two Cα atoms of these residues is below the threshold. We varied the threshold from 6 Å to 15 Å in 1 Å increments to generate an original contact map for each case. After the reconstruction, we matched the scale for each reconstruction so that the mean distance between points *i* and *i+1* for each *i* was 3.8 Å (Supplementary Fig. 2). In these examples, our reconstructions tend to be more accurate from the viewpoint of the 3D correlation coefficient (Hirata *et al.*, 2016) than that of Lesne *et al.* (2014) when the local distance is set to 1 without using the Jaccard coefficient. See Eq. (1) in the Supplementary Material for more details.

As a second test, we examined the proposed algorithm with the last step of the refinement process. Here, we used a polymer simulation of chromosomes at a 1-Mb resolution by (Qi *et al.*, 2020). We varied two parameters. The first was the threshold for defining the closeness, or the recurrence rate, which shows the ratio of intersections where contacts exist. The second was the number of points that kept contact information used for our reconstruction. During the coarse reconstruction, every fifth point was used as a reconstructed point. The reproduced contact map has a higher accuracy than that reconstructed using a portion of points (Supplementary Figs. 3 (a) and (b)). Supplementary Figs. 3(c) and 3(d) show that the values of the 3D correlation coefficients for the proposed method are 0.9 or higher, indicating that the original shape is mostly preserved after the reconstruction even if a large portion of points is discarded (Supplementary Fig. 3(d)).

A further examination of the results showed that the 3D correlation coefficient with the original shape tends to be systematically higher for our reconstructions than the simple application of Lesne *et al.* (2014) followed by refinement. This result demonstrates that our proposed framework can reconstruct finer detailed structures more effectively. This may be because the ratio of points used to estimate the local distances in the proposed method is a robust quantity under uniform sparsity.

## 3. Results

We analyzed the datasets of Tan *et al.* (2018). There are two types of cells: GM cells (GM12878), which are a female human lymphoblastoid cell line, and peripheral blood mononuclear (PBMC) cells. The datasets were downloaded from www.ncbi.nlm.nih.gov/geo/query/acc.cgi?acc=GSE117876 with the GEO Series accession number GSE117876. We used their “clean” datasets for our reconstructions. Below, we show the results of 15 GM cells and all 18 PBMC cells. One GM cell (GM cell 8) had a missing dataset. Additionally, our reconstruction was not completed for another (GM cell 10), which may be because the chromosomes are separated into two pieces or more.

### 3.1 Fractal globule and chromosome territories

Figure 2(a) shows a typical example of our reconstruction. The set of chromosomes forms a sphere. The center typically has a hole (Fig. 2(b)). We normalized the radial distance for the hole by the mean radial distance for the reconstructed points. Then we compared the value obtained for human lymphocytes from their nucleolar area (Berger, 2008) and nucleus volume (Loiko *et al.*, 2006) (Supplementary Fig. 4). GM cells are lymphoblastoid cells, which originate from lymphocytes. The values obtained for our reconstructions are close to the estimated values for human lymphocytes. Thus, we presume that this hole corresponds to the nucleolus. In addition, the values obtained for our reconstructions are more consistent with the estimated values than those for the reconstructions in Tan *et al.* (2018).

**Figure 2:**
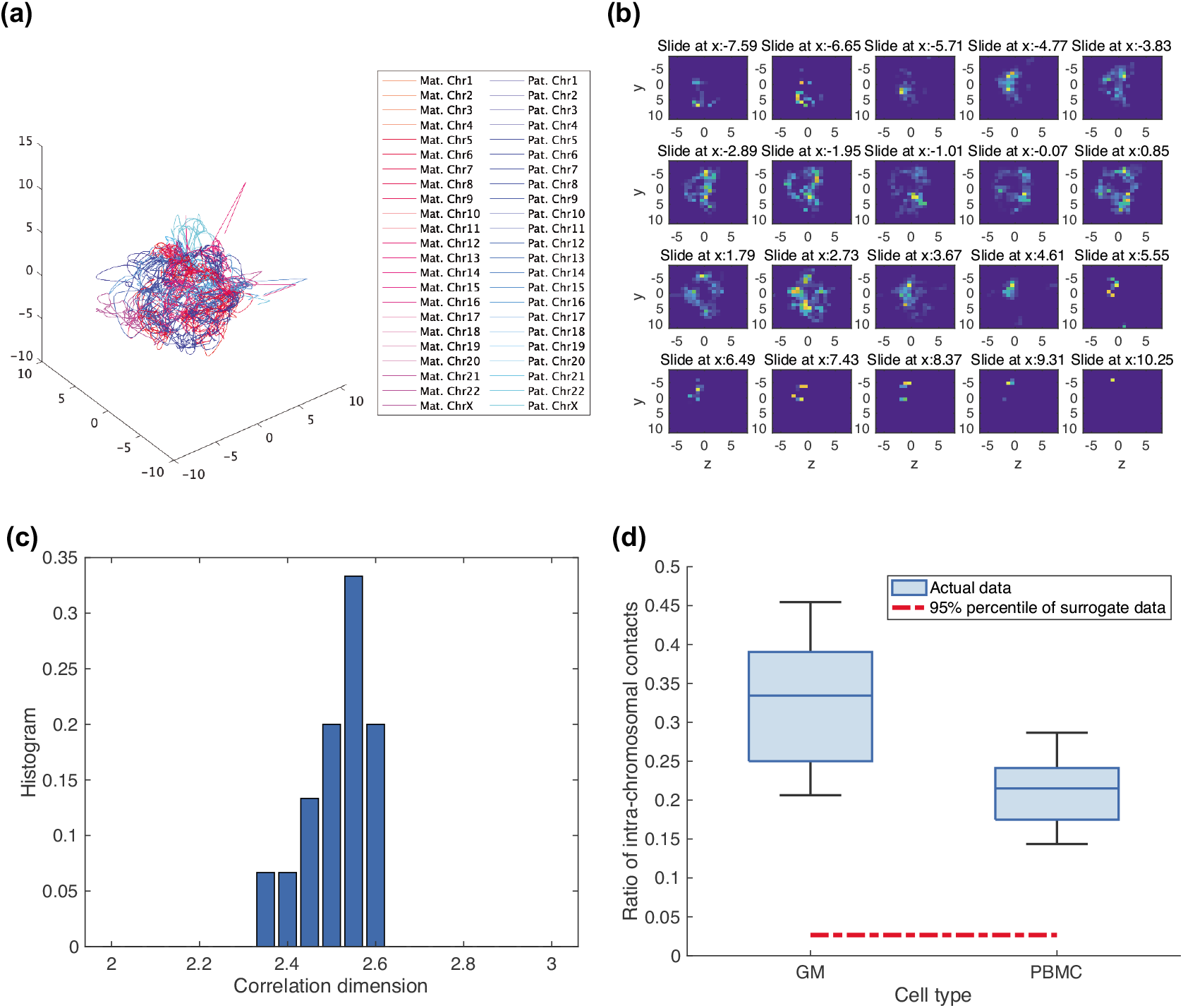
3D structure for the reconstructed chromosomes from single-cell Hi-C data. (a) Our 40-kb reconstruction for GM cell 2. (b) Density plot for GM cell 2 at a 40-kb resolution. (c) Correlation coefficients for GM cells at a 100-kb resolution. (d) Ratio of intra-chromosomal contacts for GM cells as well as PBMC cells at a 100-kb resolution compared with 20 randomly shuffled reconstructions.

When we estimated the correlation dimensions with Grassberger and Procaccia (1983), the logarithms for the accumulated proportions of the spatial distances are linearly scaled with the logarithms for the spatial distances (Supplementary Fig. 5). Moreover, the values of the correlation coefficients are between 2 and 3 (Fig. 2(c)). However, the dimension for the chromosomal structure as judged by the distribution is likely a non-integer value (average 2.5126, s.d. = 0.0745), implying a fractal chromosome structure.

It has been argued that the fractal nature of the chromosome structure leads to “chromosomal territories” (Mirny, 2011). Thus, the evaluated ratio of intra-chromosomal contacts suggests chromosomal territories rather than randomly shuffled points of chromosomes (Fig. 2(d)). Therefore, our observations imply the existence of chromosomal territories. Plotting the two alleles of each chromosome separately shows that alleles are clustered (Supplementary Fig. 6). Thus, our findings are similar to those by Tan *et al.* (2018).

### 3.2 Reconstruction consistency

Although our observations support a fractal globule forming chromosomal territories, there are differences between our reconstructions and those by Tan *et al.* (2018). Our reconstructions have a higher consistency for phased pairs and half-phased pairs than those by *Tan et al. (2018)* (Fig. 3). Additionally, Tan *et al.* (2018) has a higher value of reconstruction consistency for unphased pairs (Fig. 3). This artificially high value is due to their definition, which indicates that the shortest distance among four possible combinations is within the detection limit length. If each chromosome is evaluated, only our reconstructions in the sex chromosomes in PBMC cells are similar to those by Tan *et al.* (2018) (Supplementary Fig. 7(b)). Our reconstructions for other chromosomes differ (Supplementary Figs. 7(a) and 7(b)). In the sex chromosomes, the two reconstructions look similar because Tan *et al.* (2018) did not have to impute alleles. Hence, our reconstructions are more consistent with a given single diploid cell Hi-C dataset as the imputations in Tan *et al.* (2018) may contain bias.

**Figure 3:**
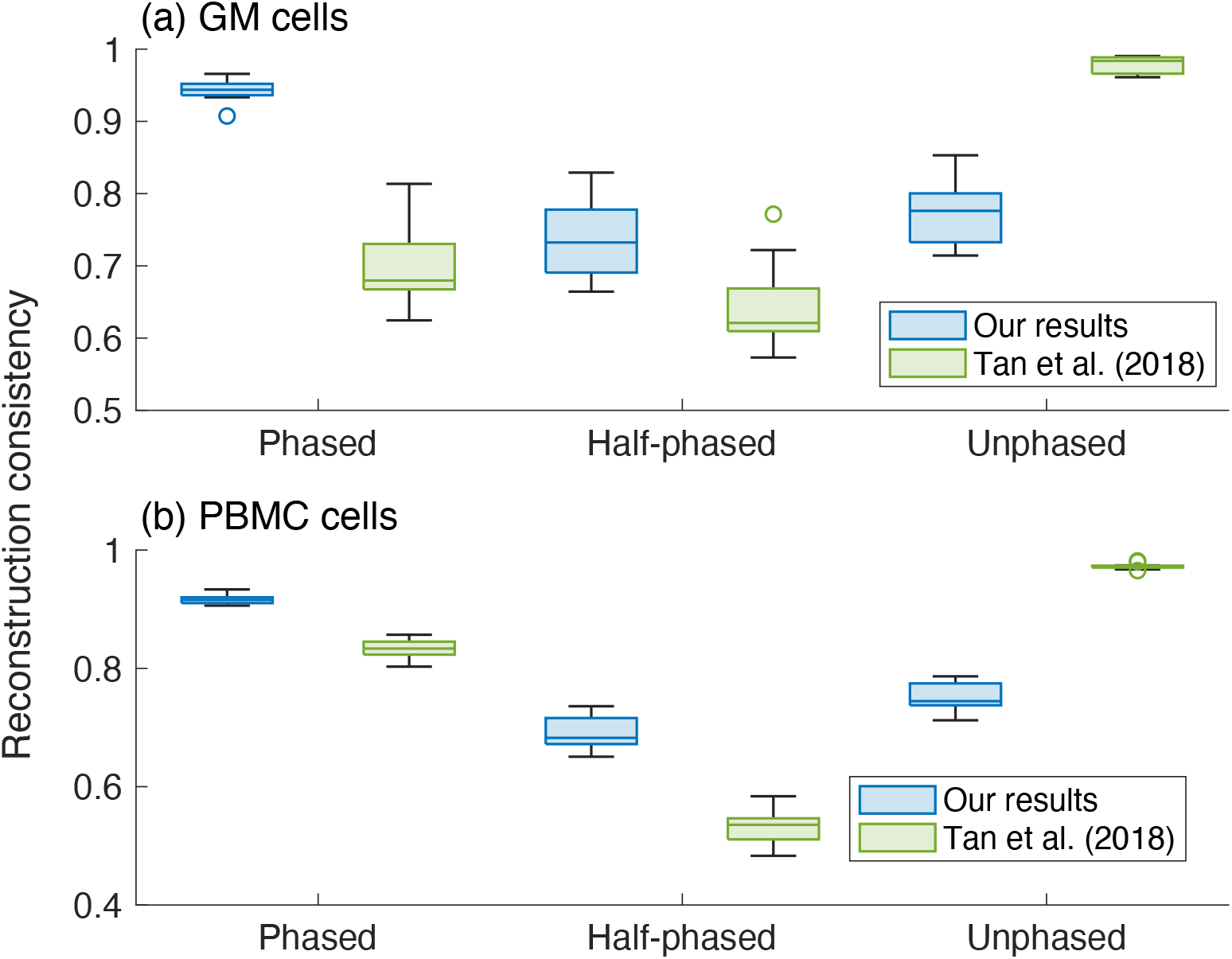
Self-checking results of the reconstructed 3D structure at a 40-kb resolution. Ratios where two close segments in a chromosomal contact for a single Hi-C are within the detection limit distance of 22/27 in the proposed method (see Supplementary Material), depending on three conditions: both segments with SNVs (phased), only one of two segments with SNVs (half-phased), and both segments without SNVs (unphased). For the results of Tan *et al.* (2018), the detection limit distance is obtained by multiplying 22/27 and the ratio of the estimated mean distance for the reconstructions of Tan *et al.* (2018) against that of our results. Panel (a) GM cells and (b) PBMC cells, where our results are on the left and those of Tan *et al.* (2018) are on the right.

Furthermore, two alleles on chromosomes 9, 13, 14, 15, 21, and 22 have similar shapes (Supplementary Fig. 7(c)). For these chromosomes, the cell-to-cell variability seems negligible (Supplementary Fig. 7(d)). Moreover, the two X chromosomes in female-derived GM cells look different in the top panel of Supplementary Fig. 7 (c). This may be due to the inactivation of one of the two X chromosomes (DISTECHE and BERLETCH, 2015).

### 3.3 Radial distance for each allele

The differences in Section 3.2 lead to the following qualitative differences. Our reconstructions reveal that one of the X chromosomes (possibly an inactive one) in female-derived GM cells is in the nuclear periphery, which is enriched with heterochromatin (Fig. 4(a)). On the other hand, the active X chromosome in male-derived PBMC male cells is closer to the center of the nucleus, which has a higher abundance of euchromatin (Fig. 4(b)). This tendency is not observed in the reconstructions by Tan *et al.* (2018) (Fig. 4(b)). In addition, the radial distances in our reconstructions for GM cells correlate well with those obtained by the FISH data for lymphoblast nuclei (Boyle *et al.*, 2001) (correlation coefficient: 0.4369). On the other hand, those for PBMC cells are not correlated with the FISH data for lymphoblast nuclei (correlation coefficient: –0.0499). This may be due to the mismatch of cell types and the different shapes of chromosomes between GM cells and PBMC cells (the middle panel of Supplementary Fig. 7(d)). In our reconstructions, chromosomes with the top five over-expressed genes in PMBC cells tend to be closer to the center of the nucleus than in GM cells (Fig. 4(d)). Such tendencies are not observed in the reconstructions by Tan *et al.* (2018) (Fig. 4(d)). Thus, our new algorithm realizes more accurate reconstructions and chromosome structures using a sparse Hi-C dataset from single diploid cells.

**Figure 4:**
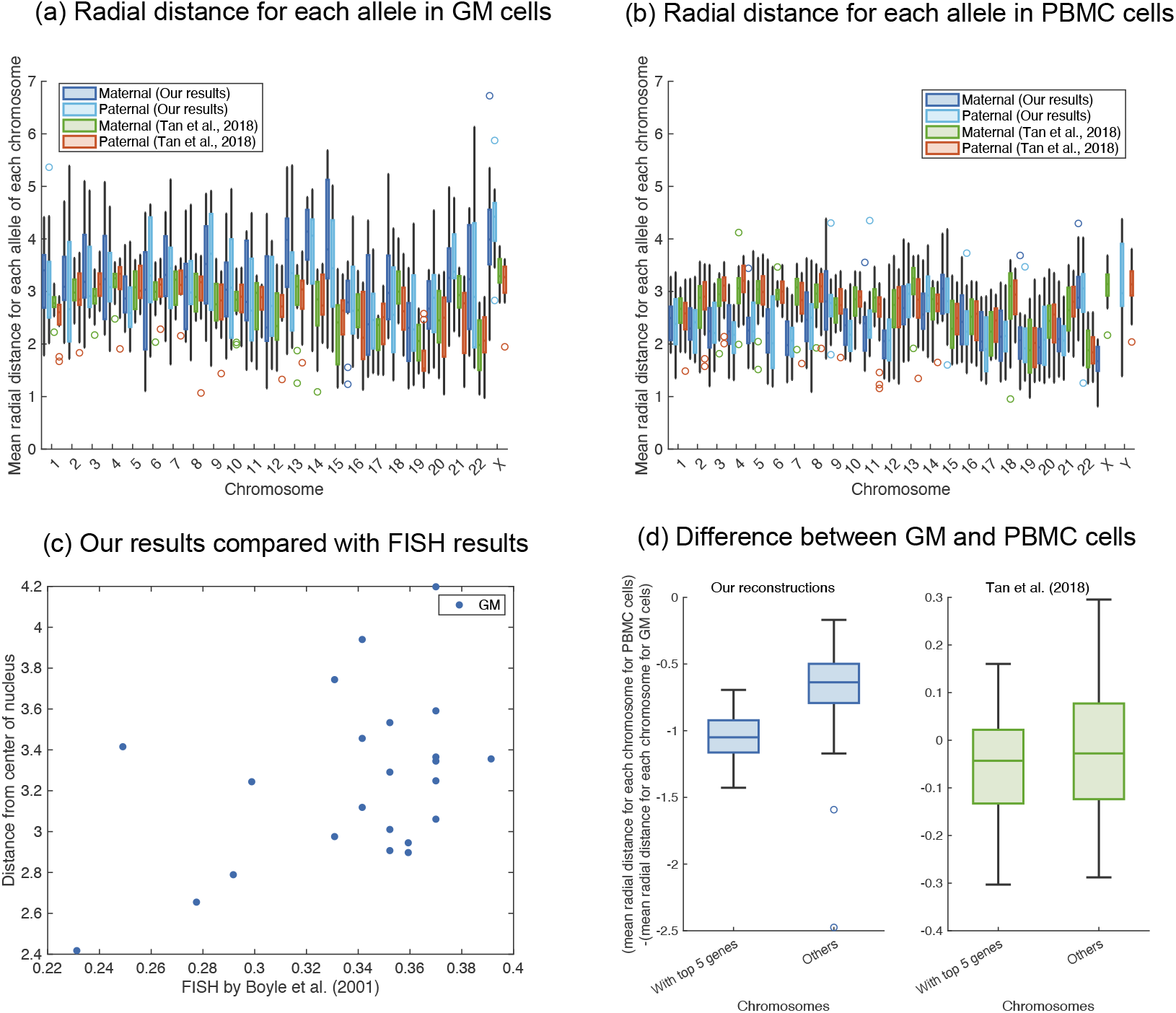
Comparisons of the reconstructed structures by radial distances for each allele at a 40-kb resolution. Panels (a) and (b) show boxplots of the radial distribution distributions for the corresponding alleles for GM cells and PBMC cells, respectively. For each chromosome, from left to right is our maternal allele, our paternal allele, the maternal allele from Tan *et al.* (2018), and the paternal allele from Tan *et al.*, (2018), except for the sex chromosomes for PBMC cell in panel (b), where the left is our reconstructions and right is those of Tan *et al.* (2018). Here, GM cells 01 and 04 are excluded because they have some chromosomes without contact information. Panel (c) shows the scatterplot of the mean radial distance over all valid cells of our GM cell reconstructions versus the FISH results in Boyle *et al.* (2001). Panel (d) highlights the differences in the radial distances between PBMC cells and GM cells in the top five genes differentially expressed in PBMC cells (Joehanes *et al.*, 2012). According to Joehanes *et al.* (2012), the top five differentially expressed genes are located on chromosomes 7, 3, 15, 4, and 14.

## 4. Discussion

These findings suggest that our reconstructions are more precise than those by Tan *et al.* (2018) for the given Hi-C datasets of single human diploid cells. The difference in accuracy may be due to the assumptions. Tan *et al.* (2018) assumed that i) two alleles are spatially separated (Supplementary Fig. 6) and ii) two alleles take different shapes (Supplementary Fig. 7(c)). On the other hand, we assumed that two consecutive points of reconstructions on the same chromosomes are close to each other.

In summary, we propose a new method to analyze a sparse Hi-C dataset of a single diploid cell with a recurrence plot-based technique. Only phased pairs are used to reconstruct the chromosome structure. Then our reconstruction is refined using an analogy of the nonlinear time series prediction. Compared to the previous reconstruction, checking the reconstruction consistency with phased pairs, half-phased pairs, and unphased pairs improves the consistency. We also demonstrate that human chromosomes take fractal shapes and form chromosomal territories. In addition, our reconstructions provide consistent results that active chromosomes are located closer to the center of the nucleus, while inactive chromosomes are located in the nuclear periphery. We hope that the proposed method will be useful to reconstruct the chromosome structure more faithfully with a given single diploid cell Hi-C dataset.

## Acknowledgements

Y.H. was partially supported by AMED (Grant Number JP21gm1310004). K.O. was supported by JST CREST, Japan (Grant Number JPMJCR18S3). M.S. was supported by JSPS KAKENHI (Grant Number JP21K12068). The funders had no role in the study design, data collection and analysis, decision to publish, or preparation of the manuscript.

